# Eilat virus (EILV) causes superinfection exclusion against West NILE virus (WNV) in a strain specific manner in *Culex tarsalis* mosquitoes

**DOI:** 10.1101/2023.05.25.542294

**Authors:** Renuka E. Joseph, Jovana Bozic, Kristine L. Werling, Nadya Urakova, Jason L. Rasgon

## Abstract

West Nile virus (WNV) is the leading cause of mosquito-borne illness in the United States. There are currently no human vaccines or therapies available for WNV, and vector control is the primary strategy used to control WNV transmission. The WNV vector Culex tarsalis is also a competent host for the insect-specific virus (ISV) Eilat virus (EILV). ISVs such as EILV can interact with and cause superinfection exclusion (SIE) against human pathogenic viruses in their shared mosquito host, altering vector competence for these pathogenic viruses. The ability to cause SIE and their host restriction make ISVs a potentially safe tool to target mosquito-borne pathogenic viruses. In the present study, we tested whether EILV causes SIE against WNV in mosquito C6/36 cells and Culex tarsalis mosquitoes. The titers of both WNV strains—WN02-1956 and NY99—were suppressed by EILV in C6/36 cells as early as 48–72 h post superinfection at both multiplicity of infections (MOIs) tested in our study. The titers of WN02-1956 at both MOIs remained suppressed in C6/36 cells, whereas those of NY99 showed some recovery towards the final timepoint. The mechanism of SIE remains unknown, but EILV was found to interfere with NY99 attachment in C6/36 cells, potentially contributing to the suppression of NY99 titers. However, EILV had no effect on the attachment of WN02-1956 or internalization of either WNV strain under superinfection conditions. In Cx. tarsalis, EILV did not affect the infection rate of either WNV strain at either timepoint. However, in mosquitoes, EILV enhanced NY99 infection titers at 3 days post superinfection, but this effect disappeared at 7 days post superinfection. In contrast, WN02-1956 infection titers were suppressed by EILV at 7 days post-superinfection. The dissemination and transmission of both WNV strains were not affected by superinfection with EILV at either timepoint. Overall, EILV caused SIE against both WNV strains in C6/36 cells; however, in Cx. tarsalis, SIE caused by EILV was strain specific potentially owing to differences in the rate of depletion of shared resources by the individual WNV strains.

**AUTHOR SUMMARY:** West Nile virus (WNV) is the main cause of mosquito-borne disease in the United States. In the absence of a human vaccine or WNV-specific antivirals, vector control is the key strategy to reduce WNV prevalence and transmission. The WNV mosquito vector, Culex tarsalis, is a competent host for the insect-specific virus Eilat virus (EILV). EILV and WNV potentially interact within the mosquito host, and EILV can be used as a safe tool to target WNV in mosquitoes. Here, we characterize the ability of EILV to cause superinfection exclusion (SIE) against two strains of WNV—WN02-1956 and NY99—in C6/36 cells and Cx. tarsalis mosquitoes. EILV suppressed both superinfecting WNV strains in C6/36 cells. However, in mosquitoes, EILV enhanced NY99 whole-body titers at 3 days post superinfection and suppressed WN02-1956 whole-body titers at 7 days post superinfection. Vector competence measures, including infection, dissemination, and transmission rates and transmission efficacy, as well as leg and saliva titers of both superinfecting WNV strains, were not affected by EILV at both timepoints. Our data show the importance of not only validating SIE in mosquito vectors but also testing multiple strains of viruses to determine the safety of this strategy as a control tool.

## INTRODUCTION

West Nile virus (WNV) is a single-stranded, positive-sense RNA virus belonging to the genus Flavivirus (family Flaviviridae) (1) ; since its first report in the USA in New York City in 1999 (2,3), WNV has expanded its geographical prevalence throughout North America (4) and is now the predominant cause of mosquito-borne disease in the USA (5). WNV is maintained in nature by an enzootic cycle between Culex mosquitoes and birds; however, spillovers into dead-end hosts such as humans and horses frequently occur, triggering epidemics (1). The severity of symptoms in humans varies from asymptomatic WNV infection to neuroinvasive disease and death (5,6). Since 1999, the USA has had 50,000 confirmed WNV cases; 25,000 neuroinvasive cases; and 2000 deaths (5,6).

Since the first WNV outbreak in the USA in 1999 WNV has evolved, with the causative WNV genotype NY99 being displaced by the genotype WN02, which currently circulates in the US population (7,8). The WN02 genotype sequence differs from the NY99 genotype sequence by three nucleotides (7,8). Two of these mutations were in the envelope protein and one in the NS5 protein, but only one of these mutations was non-synonymous (7,8). The WN02 genotype is characterized by a single amino acid change in the envelope protein from valine to alanine at position 159 (VE159A). This mutation, VE159A, enhances the vector competence of the WN02 genotype in Culex mosquitoes, particularly in Cx. tarsalis (9)—the main WNV vector in rural areas (10). Cx. pipiens and Cx. quinquefasciatus are also key WNV vectors but mainly in urban settings (11). Currently, no human vaccines or WNV-specific antivirals are available, making vector control the primary strategy used to reduce WNV transmission (12).

The WNV vector Cx. tarsalis is also a competent host for Eilat virus (EILV) (13), an insect-specific virus (ISV) belonging to the genus Alphavirus (family Togaviridae) (13). EILV is a small, enveloped, single-stranded positive-sense RNA virus that is unable to infect vertebrate cells at both the attachment/entry and replication stages of the viral life cycle (14,15). ISVs, including EILV, interact and modulate the vector competence of human pathogenic viruses in their shared mosquito hosts (16–22).A deeper understanding of these interactions can potentially be used to develop these ISVs into safe tools to target pathogenic viruses in mosquitoes.

One such interaction of interest is superinfection exclusion (SIE), a phenomenon that occurs when a pre-existing viral infection in cells blocks or interferes with a secondary infection of the same virus (homologous interference) (23–25) or closely related virus (26–28), or even an unrelated virus (heterologous interference) (21,29,30). The precise mechanism(s) of SIE remains unknown. However, there is evidence suggesting that the primary virus impacts different stages of the virus life cycle of the challenge virus in cells, including attachment (31,32), penetration (24), and replication (33,34).

ISVs cause both homologous and heterologous interference in cell culture and in mosquitoes, although the results have been variable (19,20,22,33,35–37). For example, Palm Creek virus, an insect-specific flavivirus, when superinfected with WNV in C6/36 cells, caused heterologous interference against WNV (17) but Cx. flavivirus Izabal, another insect-specific flavivirus, had no effect on WNV under similar conditions (38). Previous studies have demonstrated that EILV causes homologous and heterologous interference against other alphaviruses in C7/C10 cells and Aedes aegypti mosquitoes (22). The superinfection of EILV-infected Aedes aegypti with Chikungunya virus (CHIKV) delayed CHIKV dissemination by 3 days (22). The mosquito species Cx. tarsalis is a more competent host for EILV than Aedes aegypti (13) and therefore, we propose to investigate the ability of EILV to cause heterologous interference against the unrelated flavivirus WNV in C6/36 cells and Cx. tarsalis mosquitoes.

## MATERIALS AND METHODS

Cells and cell culture. WNV- and EILV-EILV-susceptible Aedes albopictus mosquito cell line C6/36 was propagated in Roswell Park Memorial Institute (RPMI) medium—consisting of RPMI 1640 medium (Gibco/Thermo Fisher Scientific, Waltham, MA, USA), supplemented with 10% (v/v) fetal bovine serum (FBS; Gibco/Thermo Fisher Scientific), penicillin (100 U/ml; Gibco/Thermo Fisher Scientific), and streptomycin (100 µg/ml; Gibco/Thermo Fisher Scientific)—and maintained at 28°C with no CO_2_.

Viral cDNA clone and virus propagation. <u>Eilat virus</u>: The EILV cDNA clone was obtained from the World Reference Center for Emerging Viruses and Arboviruses at the University of Texas Medical Branch (Galveston, TX, USA) and used for all experiments. The EILV cDNA clone consists of EILV strain EO329 with an enhanced green fluorescent protein (eGFP) inserted into the nsp3 hypervariable region of the viral genome (EILV-eGFP). The EILV-eGFP virus was rescued as previously described (13).

### West Nile virus

WNV strain WN02-1956 (GenBank: AY590222) stocks were originally obtained from Dr. Gregory Ebel at the Center for Vector-Borne Infectious Diseases, Colorado State University (Fort Collins, CO, USA), whereas WNV strain NY99 (GenBank: AF196835.2) stocks were acquired from Dr. Laura Kramer at the Wadsworth Center, New York State Department of Health (Albany, NY, USA). WNV strains were amplified in C6/36 cells, and the virus-containing supernatant was collected at 7 days post infection (dpi). The collected supernatant was aliquoted and stored at -80°C until use. A focus-forming assay (FFA) was used to quantify both EILV and WNV titers (described below).

### Superinfection fluorescent imaging

EILV-<u>WNV superinfection in C6/36 cells:</u> C6/36 cells were seeded in two T75 tissue culture flasks (Corning /Falcon) at a density of 6 × 10^7^ cells and incubated overnight without CO_2_ at 28°C until they reached ∼70%–80% confluency. Both T75 flasks were then washed with serum-free RPMI medium. One of the flasks was then inoculated with EILV-eGFP at a multiplicity of infection (MOI) of 10, whereas the other flask was mock inoculated with serum-free RPMI medium; the flasks were incubated at 28°C with no CO_2_ for 1 h. After incubation, the remaining inoculum in the flasks were removed and replaced with complete RPMI, and the flasks were incubated at 28°C with no CO_2_ for 48 h.

The EILV-eGFP and mock-infected C6/36 cells were detached using trypsin (Gibco/Thermo Fisher Scientific) after the 48 h incubation and seeded in a 24-well tissue culture plate (Greiner Bio-One) at a density of 5 × 10^6^ cells in triplicate and incubated overnight at 28°C with no CO_2_. The cells in both groups were washed with serum-free RPMI medium, infected with the WNV strain WN02-1956 or NY99 at MOIs of 0.1 or 0.01, and incubated at 28°C with no CO_2_ for 1 h. The cells were washed twice with complete RPMI medium after WNV infection to remove any unattached virus. Complete RPMI medium was then added to the cells, and the EILV–WN02-1956-superinfected or EILV–NY99-superinfected and the control WN02-1956-infected or NY99-infected cells at both MOIs were incubated at 28°C with no CO_2_ for 72 h.

Superinfected cells were fixed 72 h post superinfection using 300 µl of 4% formaldehyde in 1× phosphate-buffered saline (PBS) for 30 min at room temperature (RT). Superinfected cells were washed twice with 500 µl of 1× PBS and then permeabilized by adding 300 µl of 0.2% triton-X in 1× PBS to each well for 15 min at RT. Then, the cells were washed twice with 1× PBS and blocked with 300 µl of 3% bovine serum albumin (BSA) in 1× PBS for 1 h at RT. The cells were washed again with 1× PBS twice; next, 250 µl of the primary antibody, monoclonal anti-flavivirus group antigen antibody (clone D1-4G2-4-15, diluted 1:500 in 3% BSA), was added to each well, and the cells were incubated overnight at 4°C. Superinfected cells were then washed twice with 1× PBS, and 250 µl of the secondary antibody, goat anti-mouse IgG (H+L) highly cross-adsorbed secondary antibody (Alexa Fluor Plus 594, diluted 1:1000 in 3% BSA), was added to each well. The plates were wrapped in aluminum foil and incubated overnight at 4°C. Finally, cells were washed twice and maintained in 300 µl of 1× PBS to prevent drying. EILV infection was observed under the fluorescein isothiocyanate (FITC) filter and WN02-1956 and NY99 infections under the tetramethylrhodamine isothiocyanate (TRITC) filter on the ECHO Revolve microscope. FITC and TRITC images were merged (all scale bars correspond to 180 µm).

### In vitro superinfection assay

EILV-eGFP-infected C6/36 cells were superinfected with WN02-1956 or NY99 at MOIs 0.1 and 0.01 in 24-well plates as described above. Superinfected and control WN02-1956-infected or NY99-infected cells at both MOIs were maintained at 28°C with no CO_2_ for 96 h. At 12, 24, 48, 72, and 96 h post superinfection, 300 µl of supernatant was collected from each superinfection and control well and replaced with fresh medium. Samples were stored at -80°C until use, and WNV titers in the samples were quantified using FFA (described below). The in vitro superinfection assay was performed in duplicate using aliquots of the same EILV-eGFP and WNV stocks that underwent the same number of freeze-thaw cycles.

### Viral attachment and internalization assay

The viral attachment and internalization assay was performed as previously described with some modifications (25). For the viral attachment assay, a 24-well tissue culture plate with EILV-eGFP-infected (MOI 10) and mock-infected C6/36 cells seeded in duplicate was superinfected with WWN02-1956 or NY99 at an MOI of 0.1 as described above. Superinfected cells were then incubated at 4°C for 1 h to allow virus particles to attach but not enter the cells. Then, superinfected cells were washed with 1x PBS (Gibco/Thermo Fisher Scientific) and then washed twice with complete RPMI medium to remove any unattached WNV. Both groups of superinfected cells, with and without EILV-eGFP infection, were resuspended in 300 µl of TRIzol for RNA extraction.

Similarly, for the viral internalization assay, a 24-well plate with EILV-eGFP-infected and mock-infected C6/36 cells superinfected with WN02-1956 or NY99 at an MOI of 0.1 was incubated at 28°C with no CO_2_ for 1.5 h to allow attachment and internalization of WNV. Superinfected cells were then washed with 1x PBS followed by a high-salt buffer containing 50 mM Na_2_CO_3_ (pH 9.5; Fisher chemicals, Thermo Fisher Scientific) and 1 M NaCl (Fisher chemicals, Thermo Fisher Scientific) for 3 min at RT. The high-salt buffer was removed, and the superinfected cells were resuspended in 300 µl of TRIzol for RNA extraction. RNA was extracted from all samples using the phenol/chloroform extraction method as previously described (39) and DNA contamination in the extracted RNA was removed from the samples using the TURBO DNA-free kit (Invitrogen, Thermo Fisher Scientific). Total RNA was quantified using NanoDrop ND-1000 (NanoDrop Technologies/Thermo Fisher Scientific), and WNV RNA in 60 ng of total RNA from each sample was quantified by reverse transcription-quantitative polymerase chain reaction (RT-qPCR; described below). The virus attachment and internalization assays were performed in duplicate using aliquots of the same EILV-eGFP and WNV stocks that underwent the same number of freeze-thaw cycles.

### Reverse transcription-quantitative polymerase chain reaction (RT-qPCR)

WNV RNA was quantified using the Qiagen rotor gene Q-compatible qScript One-Step SYBR Green RT-qPCR kit (Quantabio). The RT-qPCR reaction was set up as per the manufacturer’s instructions, using WNV envelope protein-specific primers 51-TTGCAAAGTTCCTATCTCGTCAG-31 and 51-ACATGCCTCCGAACAGTGAG-31 for both reverse transcription and amplification of the target WNV sequence. The WNV polyprotein DNA fragment (1917–2338 bp) synthesized by IDT technologies served as standards for the RT-qPCR reaction, making a standard curve from 10^6^ to 10^3^ viral copies/µl. All samples and standards were run in duplicate, and WNV titers were determined using the standard curve.

### Mosquitoes and mosquito rearing

EILV-eGFP and WNV-competent mosquitoes, Cx. tarsalis (KNWR strain), were used in all experiments in our study. The KNWR strain was obtained from Dr. Christopher Barker at the University of California–Davis School of Veterinary Medicine (Davis, CA, USA). Uninfected mosquitoes were reared and maintained at the Millennium Sciences Complex (The Pennsylvania State University, University Park, PA, USA), whereas WNV-infected mosquitoes were maintained at the Eva J. Pell Biosafety Level 3 (BSL3) laboratory (The Pennsylvania State University, University Park, PA, USA), as previously described (13). Cx. tarsalis KNWR larvae were reared in 30 × 30 × 30 cm cages, and growth conditions for both KNWR adults and larvae were 25°C ± 1°C, 16:8 h light: dark diurnal cycle, and 80% relative humidity. Larvae were fed TetraMin (Tetra), and all adult mosquitoes were fed with 10% sucrose solution-soaked cotton balls ad libitum.

### Mosquito superinfection assay

Adult Cx. tarsalis KNWR female mosquitoes (3–5 days post emergence) were sugar-starved for 24 h and then fed an infectious blood meal consisting of 1:1 anonymous human blood (BioIVT) and 10^7^ FFU/ml of EILV-eGFP (EILV-eGFP-infected group) or RPMI medium (mock-infected group) at 37°C using a water-jacketed membrane feeder. Previous studies have shown that an EILV-eGFP dose of 10^7^ FFU/ml leads to a robust EILV infection in ∼70% of the blood-fed Cx. tarsalis KNWR mosquitoes (13). After the infectious bloodmeal, mosquitoes from both groups were cold-anesthetized, and fully engorged female mosquitoes were counted and sorted into different cardboard cup cages with 10% sucrose solution-soaked cotton balls until further experimentation. At 5 dpi, mosquitoes in both groups were allowed to lay eggs in plastic cups filled with deionized water and the eggs were then discarded. Both EILV-eGFP-infected and mock-infected mosquitoes were moved to the Eva J. Pell BSL3 laboratory at 6 dpi and sugar-starved for 24 h. Mosquitoes in both groups were then fed again at 7 dpi with an infectious blood meal containing 1:1 anonymous human blood and WN02-1956 or NY99 at a dose of 10^7^ FFU/ml. Fully engorged females from the superinfected and single-infected control groups were cold-anesthetized and placed into cardboard cup cages with 10% sucrose solution-soaked cotton balls. These cages with WNV-infected mosquitoes were then double caged and maintained at 25°C ± 1°C, 16:8 light: dark diurnal cycle, and 80% relative humidity until assay timepoints at 3 and 7 days post superinfection.

Whole bodies from both superinfected and singly infected control groups were collected 3 days post superinfection and placed into 2 ml microcentrifuge tubes containing 300 µl mosquito diluent (20% FBS, 100 µg/ml of streptomycin, 100 µg /ml of penicillin, 50 µg/ml gentamicin [Gibco/Thermo Fisher Scientific], and 2.5 µg/ml amphotericin B [Gibco/Thermo Fisher Scientific] mixed in 1× PBS). Samples were then homogenized using a battery-operated tissue homogenizer (VWR) and disposable pestles (Fisher scientific/Thermo Fisher Scientific). Similarly, at 7 days post superinfection, mosquitoes from both groups were forced to salivate for 30 min at RT into a capillary glass tube containing a mixture of 1:1 FBS and 50% sucrose. After salivation, the saliva was expelled into a 2 ml microcentrifuge tube containing 100 µl mosquito diluent. Legs and bodies of superinfected and control mosquitoes were then collected in 2 ml microcentrifuge tubes containing 300 µl mosquito dilutant. The legs and bodies were homogenized using the battery-operated homogenizer and disposable pestles, and all samples were stored at -80°C until further processing. The WNV titers of all samples collected were quantified using FFA (described below). The mosquito superinfection assay was performed in duplicate, and aliquots of the same EILV-eGFP and WNV stocks that had undergone the same number of freeze-thaw cycles were used in both replicates.

Using the FFA results, the following vector competence parameters were determined. The infection rate (IR) was calculated as the proportion of WNV-infected mosquitoes among the total number of EILV-eGFP or mock-infected mosquitoes; the dissemination rate (DIR), as the proportion of WNV-infected mosquitoes with WNV-positive legs; the transmission rate (TR), as the proportion of mosquitoes with WNV-positive saliva from those with WNV-positive legs; and the transmission efficacy (TE), as the proportion of mosquitoes with WNV-positive saliva among the total number of EILV-eGFP or mock-infected mosquitoes.

### Focus-forming assay

We quantified EILV-eGFP titers using FFA as previously described (13). Similarly, WNV titers were quantified by FFA as previously described but with some modifications (21). 96-well plates were seeded with C6/36 cells at a density of 1 × 10^5^ cells/well and incubated at 28°C with no CO_2_ overnight. The complete RPMI medium from the wells was then removed. Samples from the in vitro and in vivo superinfection assays were then serially diluted in serum-free RPMI medium from 10^0^ to 10^-7^ and 10^0^ to 10^-3^, respectively, and 30 µl of each diluted sample was added to the prepared C6/36 cells in duplicates. Mosquito saliva samples were not diluted, and 30 µl of each sample was added directly to the prepared cells. The cells were then incubated at 28°C with no CO_2_ for 1 h, after which the samples were removed, and the cells were covered with 100 µl of RPMI containing 0.8% methylcellulose (Sigma-Aldrich). The WNV-infected cells were incubated at 28°C without CO_2_ for 48 h. At 48 h post infection, the infected C6/36 cells were fixed using 50 µl of 4% formaldehyde (Sigma-Aldrich) in 1× PBS for 30 min at RT. Cells were washed twice with 100 µl 1× PBS and were then permeabilized with 30 µl of 0.2% triton-X in 1× PBS for 15 min at RT. The washing step was repeated, and the cells were blocked with 30 µl of 3% BSA in 1× PBS for 1 h at RT. The cells were washed again, and 30 µl of the primary antibody, monoclonal anti-flavivirus group antigen antibody (clone D1-4G2-4-15 [BEI resources], diluted 1:500 in 3% BSA), was added to each well. The cells were then incubated overnight at 4°C. Cells were then washed to remove any unattached primary antibody, and 30 µl of the secondary antibody goat anti-mouse IgG (H+L) highly cross-adsorbed secondary antibody (Alexa Fluor Plus 594 diluted 1:1000 in 3% BSA) was added to each well. The plates were wrapped in aluminum foil and incubated overnight at 4°C. After a final washing step, the cells were maintained in 100 µl of 1× PBS to prevent drying. Fluorescent foci of EILV-eGFP were counted using the FITC filter on the ECHO Revolve microscope, whereas WN02-1956 or NY99 foci were counted using the TRITC filter.

### Statistical analysis

A two-way ANOVA followed by the Tukey test was used to determine the difference between WNV replication kinetics under the superinfection and single infection conditions in C6/36 cells. The WNV viral titers from the post-attachment or internalization assay were compared using Mann-Whitney U tests. The WNV viral titers of the body, leg, and saliva samples from superinfected and singly infected Cx. tarsalis were also compared using Mann-Whitney U tests. The differences in the WNV IR, DIR, TR, and TE post superinfection in EILV-infected or mock-infected Cx. tarsalis mosquitoes were evaluated using Fisher’s exact test. GraphPad Prism version 9.0.4 was used to perform all statistical tests.

## RESULTS

Eilat virus caused heterologous interference against West Nile virus strains WN02-1956 and NY99 in vitro. To investigate the ability of EILV to cause SIE against WNV in cell culture, C6/36 cells were mock infected or infected with EILV-eGFP at an MOI of 10. After EILV infection was established at 72 h post infection, cells were superinfected with the WNV strain WN02-1956 or NY99 at MOIs of 0.1 and 0.01. Superinfected cells at both MOIs were also observed using fluorescent microscopy at 72 h post superinfection to determine whether EILV-eGFP-infected cells were resistant to WNV infection. Moreover, differences in WN02-1956 or NY99 replication kinetics under superinfection and single infection conditions at 12, 24, 48, 72, and 96 h post superinfection were determined to detect SIE in C6/36 cells.

We observed that C6/36 cells infected with EILV-eGFP, determined by eGFP expression by fluorescent microscopy, were susceptible to superinfection by WN02-1956 or NY99 virus, as indicated by Alexa Flour 594 fluorescence at both MOIs tested (Fig 1A). The viruses also co-localized in the cells, suggesting co-infection by EILV and WN02-1956 or NY99 (Fig 1A).

**Figure 1.**
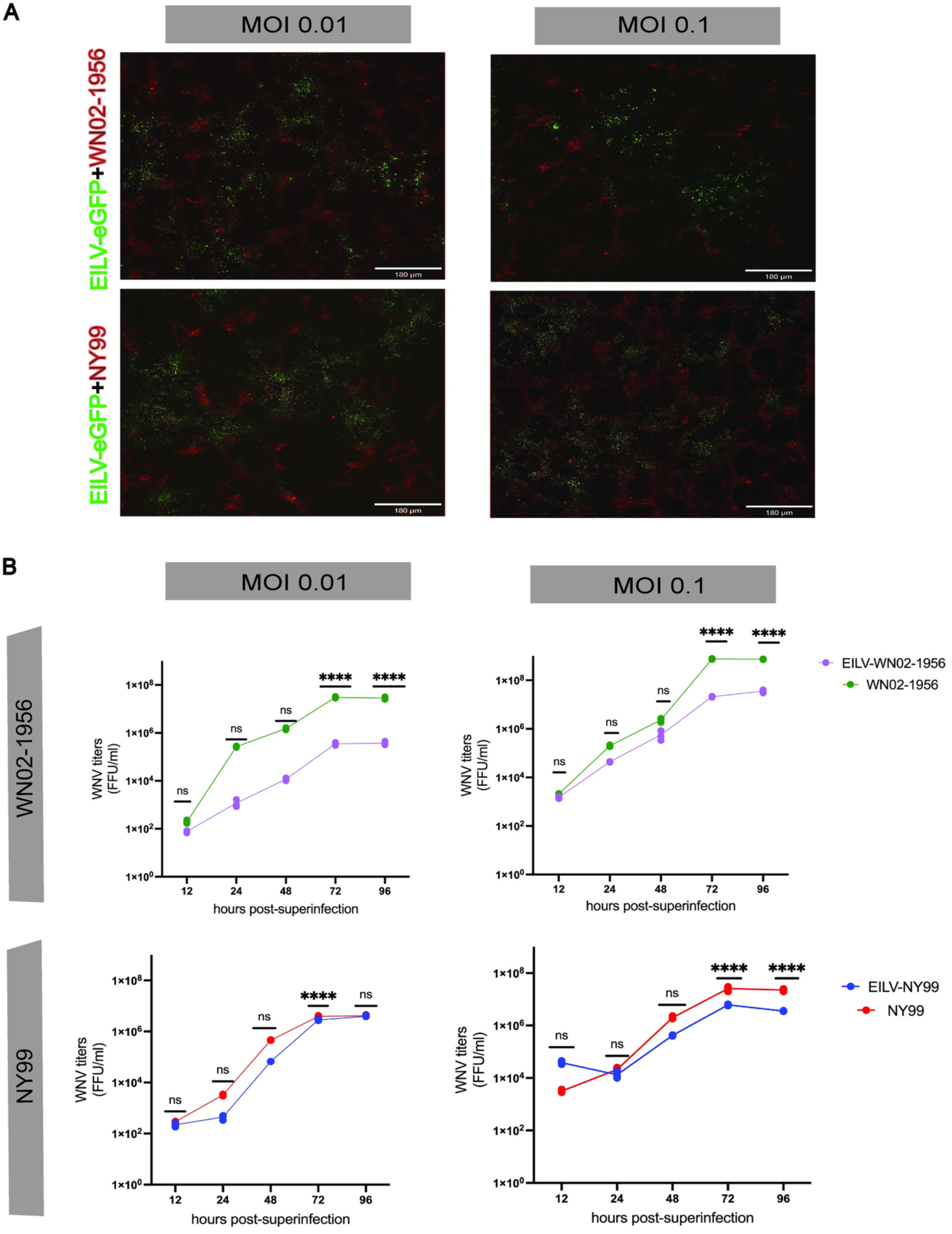
Superinfection exclusion (SIE) against WNV strains, WN02-1956 and NY99 in C6/36 cells by EILV. (A) Superinfection of EILV-eGFP-infected C6/36 cells (multiplicity of infection [MOI] of 10) with WN02-1956 or NY99 at an MOI of 0.01 and 0.1 was visualized by fluorescent microscopy. Representative images show eGFP fluorescence for EILV-eGFP and Alexa Fluor 594 fluorescence for both WNV strains. All scale bars equal 180 µm. (B) Titers of WNV at 12, 24, 48, 72 and 96 h post-superinfection in EILV-eGFP-infected C6/36 cells (MOI 10) and mock-infected cells superinfected with WN02-1956 or NY99 at an MOI of 0.01 and 0.1 were determined by focus forming assay (FFA). Each point represents a single infection at each timepoint. Statistical significance was evaluated using two-way ANOVA followed by Tukey test. ** P< 0.01 and **** P< 0.0001.

we also found that the titers of WN02-1956, were statistically significantly lower at 72 and 96 h post superinfection in EILV-eGFP-infected C6/36 cells than in mock-infected cells at both MOIs tested (Fig 1B, two-way ANOVA with Tukey test, all P < 0.0001). A similar trend was observed in the second replicate of this assay, but with a significant suppression of WN02-1956 titers occurring earlier at 48 h in superinfected cells which continued at 72 and 96 h post superinfection at both MOI 0.1 (Fig S1A; all P < 0.0001) and MOI 0.01 (Fig S1A; P < 0.0001, P < 0.0001, and P ≤ 0.01 respectively).

In contrast, the titer of the WNV strain NY99 was significantly lower in superinfected cells at MOI 0.01 than in singly infected control cells only at 72 h post superinfection (Fig 1B; P < 0.0001), with the NY99 titers rebounding at 96 h post superinfection (Fig 1B, P > 0.05). The second replicate of the EILV–NY99 superinfection assay at MOI 0.01 in C6/36 cells differed from the first replicate, with significantly lower NY99 titers observed at 48, 72, and 96 h post superinfection (Fig S1A; all P < 0.0001). NY99 titers showed some but not significant recovery at 96 h post superinfection, unlike that observed in the first replicate. Similarly, superinfection of EILV-eGFP-infected C6/36 cells with NY99 at MOI 0.1 led to significantly lower NY99 titers at 72 and 96 h post superinfection (Fig S1A, all P < 0.0001) in superinfected cells than in singly infected control cells in both replicates. Overall, the WN02-1956 and NY99 titer reduction by EILV in C6/36 cells under superinfection conditions ranged between 10-fold and 100-fold.

Eilat virus interfered with the attachment of the superinfecting virus NY99 in C6/36 cells. Having established that EILV causes SIE against WN02-1956 and NY99 in C6/36 cells, we next investigated how EILV causes heterologous interference against unrelated WNV in cells. To this end, we assessed whether EILV interferes with the attachment or internalization stage of the WN02-1956 or NY99 viral life cycle in C6/36 cells. EILV-eGFP-infected and mock-infected C6/36 cells were superinfected with WN02-1956 or NY99 at an MOI of 0.1. WNV particles were either allowed to only attach to their receptors or also internalize into the cells. WN02-1956 and NY99 RNA from superinfected and singly infected cells post attachment or internalization were quantified by RT-qPCR to determine whether EILV interferes with either of these stages of the WNV life cycle.

The viral titers of attached or internalized WN02-1956 in EILV-eGFP-infected C6/36 cells and in mock-infected C6/36 cells were not significantly different (Fig 2A and 2B; respectively, Mann-Whitney U tests, both P > 0.05). However, viral titers of attached NY99 were significantly lower in superinfected C6/36 cells than in singly infected cells (Fig 2A; P < 0.05). This difference between NY99 viral titers was not observed when comparing internalized virus particles in superinfected and singly infected cells (Fig 2B; P > 0.05).

**Figure 2.**
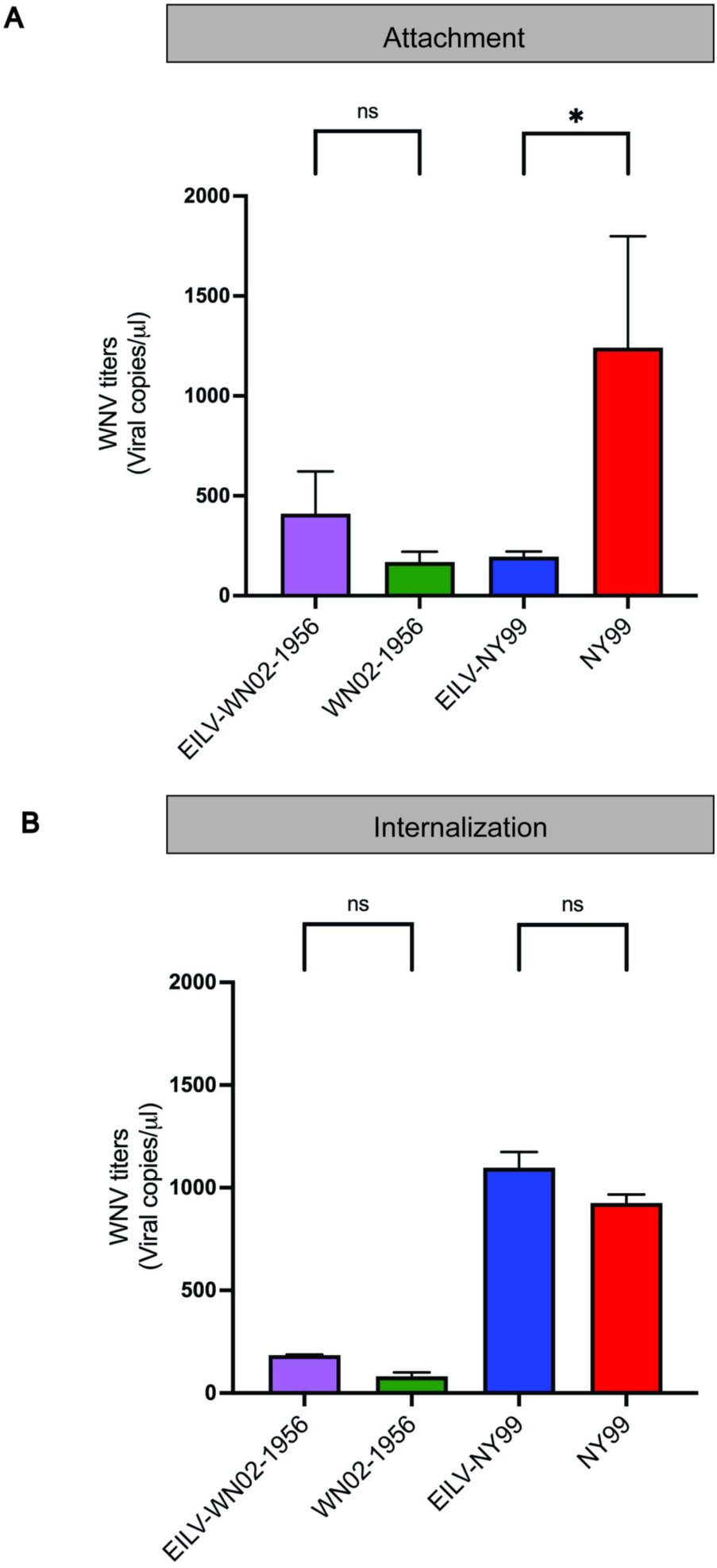
WNV attachment and internalization post-superinfection in EILV-eGFP-infected C6/36 cells. WNV titers at (A) attachment and (B) internalization stages of WNV life cycle in EILV-eGFP infected and mock-infected C6/36 cells superinfected with WN02-1956 or NY99 (MOI 0.1) were determined by RT-qPCR. Each vertical bar reflects the mean of duplicate infections, and error bars indicate standard deviation between replicates. Statistical significance was assessed using Mann-Whitney U tests. * P<0.05.

Eilat virus enhanced NY99 whole-body titers at an early timepoint post superinfection in Culex tarsalis mosquitoes. We tested whether EILV causes heterologous interference against WNV strains in mosquitoes, as previously observed in C6/36 cells, by orally superinfecting EILV-eGFP-infected Cx. tarsalis KNWR mosquitoes with WN02-1956 or NY99. We examined the presence of EILV and WN02-1956 or NY99 using FFA in the bodies of 34 and 22 EILV-eGFP-challenged KNWR mosquitoes orally challenged with 10^7^ FFU/ml of WN02-1956 or NY99, respectively, at 3 days post superinfection. Simultaneously, for our controls, we examined the presence of WN02-1956 or NY99 using FFA at 3 days post superinfection in the bodies of 29 and 19 mock-infected KNWR mosquitoes orally challenged with 10^7^ FFU/ml of WN02-1956 or NY99, respectively. The IR for mosquitoes in our superinfected or mock-infected control groups was defined as the proportion of mosquitoes with WNV infection among the total number of EILV-infected or mock-infected mosquitoes or fully engorged females, respectively.

We observed that EILV-infected Cx. tarsalis were not refractory to a secondary WN02-1956 or NY99 infection at 3 days post superinfection (Table 1). At this early timepoint, the IR of WN02-1956- and NY99-superinfected mosquitoes did not significantly differ from the IR of mosquitoes singly infected with WN02-1956 or NY99 (Table 1; Fisher’s exact tests, both P > 0.05). However, NY99 whole-body titers were significantly higher in EILV-infected mosquitoes than in mock-infected mosquitoes at 3 dpi (Fig 3B; Mann-Whitney U tests, P ≤ 0.001). No significant difference in WN02-1956 whole-body titers was observed at this timepoint between the two groups (Fig 3A, P > 0.05).

**Table 1.** Infection rate (IR) of Eilat virus (EILV)-enhanced green fluorescent protein (eGFP)-infected or mock-infected Culex tarsalis mosquitoes orally challenged with the West Nile virus (WNV) strain WN02-1956 or NY99

|  | 3 days post superinfection |
| --- | --- |
| Superinfected mosquito group | IR (%)<br>($n_I/n_T$) |
| EILV-eGFP//WN02-1956 | 53.3<br>(8/15) |
| Mock-infected/WN02-1956 | 50<br>(11/22) |
| EILV-eGFP//NY99 | 92.8<br>(13/14) |
| Mock-infected/NY99 | 89.5<br>(17/19) |

$n_I$ , number of mosquitoes infected with WNV
$n_T$ , total number of EILV-eGFP positive or mock-infected mosquitoes tested

**Figure 3.**
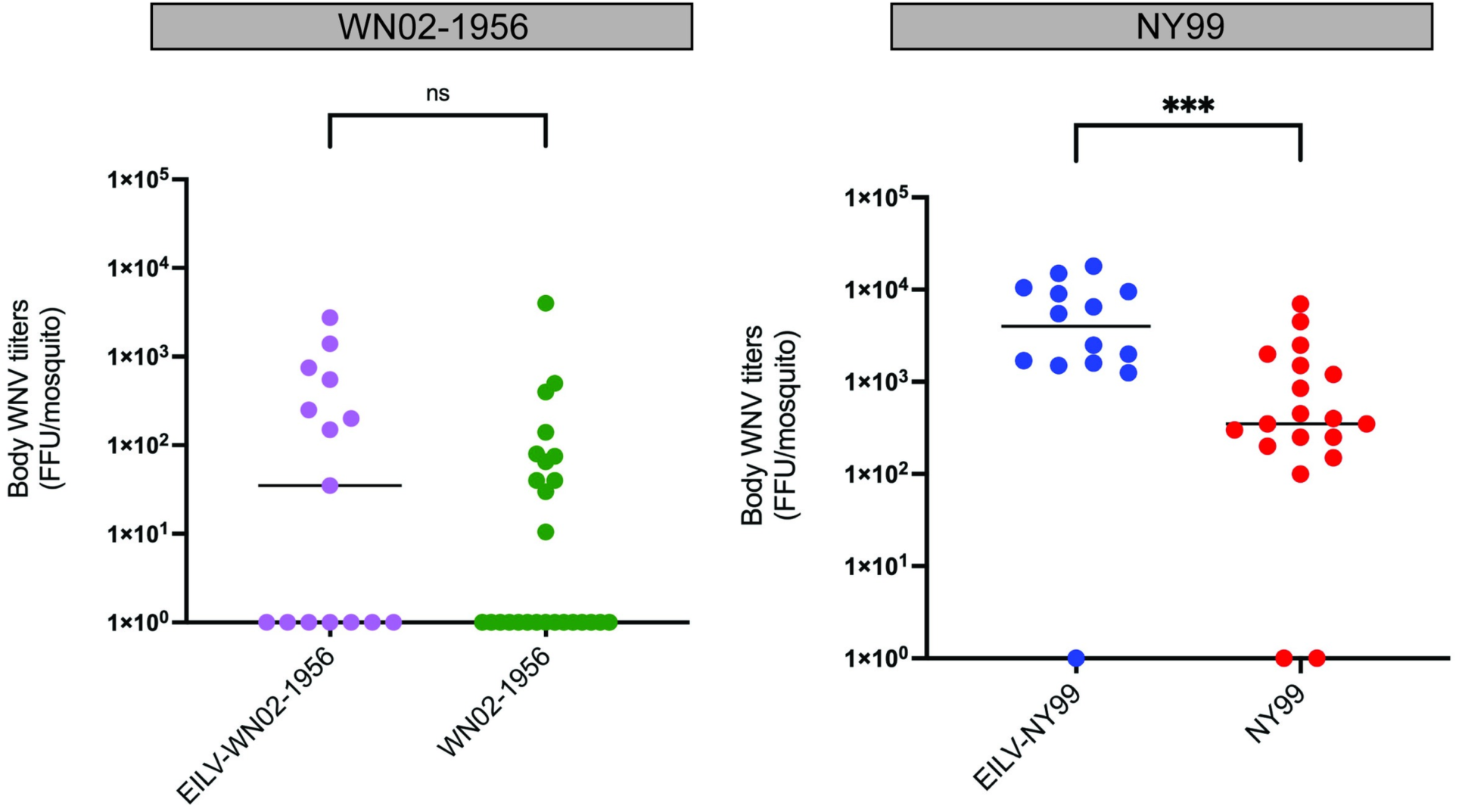
Whole-body WNV titers of superinfected and singly infected Culex tarsalis mosquitoes at 3 days post-superinfection. WNV Viral titers of whole-body samples from EILV-eGFP-infected (10^7^ FFU/ml) and mock infected Cx. tarsalis mosquitoes orally challenged with A) WN02-1956 and (B) NY99 (both 10^7^ FFU/ml) are plotted 3 days post-superinfection. Individual points represent a single mosquito sample, while group medians are depicted by horizontal bars. Significance was determined using Mann-Whitney U tests. *** P<0.001.

### Eilat virus suppressed WN02-1956 body titers at 7 days post superinfection in Culex tarsalis mosquitoes

We assessed the vector competence of WNV strains in EILV-infected and mock-infected Cx. tarsalis KNWR mosquitoes at a later timepoint to determine whether the ability of EILV to enhance or cause SIE changed with increasing time post superinfection. We determined the IR, DIR, TR, and TE of WN02-1956 in 37 EILV-infected and 36 mock-infected mosquitoes and of NY99 in 14 EILV-infected and 24 mock-infected mosquitoes at 7 days post superinfection. IR was calculated as previously defined; DIR was determined as the proportion of EILV- or mock-infected mosquitoes with WNV-positive legs, TR as the proportion of EILV- or mock-infected mosquitoes with WNV-positive saliva, and TE as the proportion of EILV-or mock-infected mosquitoes with WNV-positive saliva.

We found that the IR and DIR of WN02-1956 and NY99 in superinfected mosquitoes did not differ significantly from mock-infected mosquitoes at this timepoint (Table 2; Fisher’s exact test, P > 0.05). Both WNV strains disseminated past the midgut in EILV- and mock-infected mosquitoes, but WN02-1956 and NY99 were detected only in the saliva of mock-infected mosquitoes (Table 2, P > 0.05). However, the difference in the TR and TE of WN02-1956 and NY99 in EILV- and mock-infected mosquitoes was not statistically significant (Table 2, P > 0.05).

**Table 2.** Infection rate (IR), dissemination rate (DIR), and transmission rate (TR) and transmission efficiency (TE) of Eilat virus (EILV)–enhanced green fluorescent protein (eGFP)-positive or mock-infected Culex tarsalis mosquito species orally challenged with the West Nile virus (WNV) strain WN02-1956 or NY99

| Superinfected mosquito group | 7 days post superinfection |  |  |  |
| --- | --- | --- | --- | --- |
| | IR (%)<br>($n_I/n_T$) | DIR (%)<br>($n_L/n_I$) | TR (%)<br>($n_S/n_L$) | TE (%)<br>($n_S/n_T$) |
| EILV-eGFP/WN02-1956 | 84.2<br>(16/19) | 25<br>(4/16) | 0<br>(0/4) | 0<br>(0/19) |
| Mock-infected–WN02-1956 | 87.5<br>(28/32) | 28.6<br>(8/28) | 12.5<br>(1/8) | 3.1<br>(1/32) |
| EILV-eGFP/NY99 | 100<br>(14/14) | 28.6<br>(4/14) | 0<br>(0/4) | 0<br>(0/14) |
| Mock-infected–NY99 | 100<br>(24/24) | 50<br>(12/24) | 8.3<br>(1/12) | 4.2<br>(1/24) |

$n_I$ , number of mosquitoes infected with WNV
$n_T$ , total number of EILV–eGFP-positive or mock-infected mosquitoes tested
$n_L$ , number of mosquitoes with WNV-positive legs
$n_S$ , number of mosquitoes with WNV-positive saliva

Moreover, we found that EILV significantly suppressed WN02-1956 body titers in superinfected mosquitoes (Fig 4, Mann-Whitney U tests, P ≤ 0.001). This suppression of WN02-1956 body titers did not result in an overall decrease in WN02-1956 titers in the legs and saliva of superinfected mosquitoes (Fig 4A, P > 0.05). Similarly, no difference was observed in the NY99 body, leg, or saliva titers between superinfected and singly infected mosquitoes (Fig 4B, P > 0.05).

**Figure 4.**
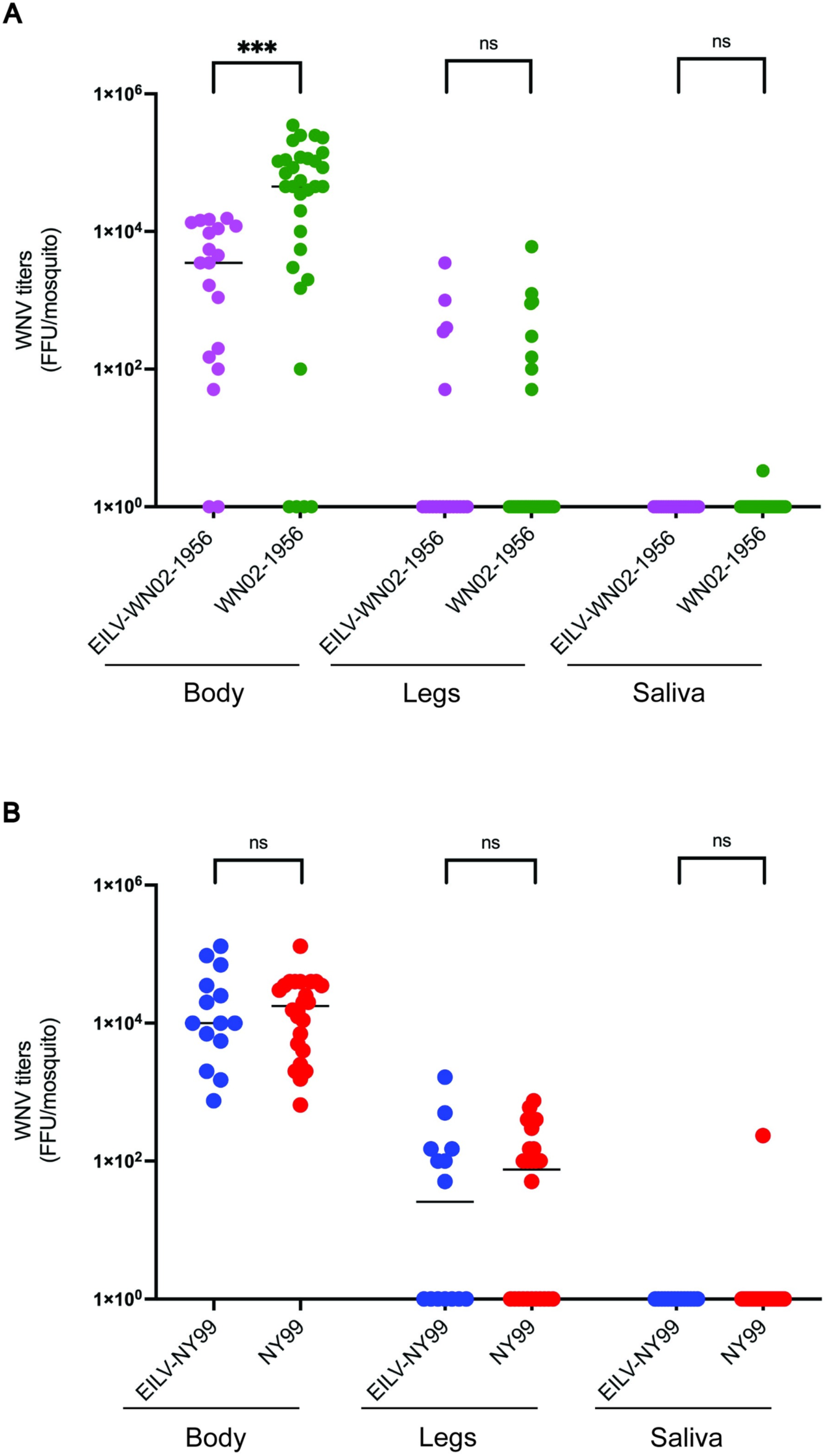
WNV titers of body, leg, and saliva samples from EILV-eGFP-infected and mock infected Culex tarsalis mosquitoes challenged with WN02-1956 or NY99. WNV titers in (A) body, (B) leg and (C) saliva samples from EILV-eGFP-infected (10^7^ FFU/ml) and mock infected Culex tarsalis mosquitoes challenged with WN02-1956 or NY99 (10^7^ FFU/ml) at 7 days post-superinfection are plotted. Individual points represent a single mosquito sample, while group medians are depicted by horizontal bars. Significance was determined using Mann-Whitney U tests. *** P<0.001.

## DISCUSSION

WNV is a pathogenic arbovirus of public health importance, especially in the USA; however, there are currently no human vaccines or WNV-specific antivirals available against this pathogen (5,12). The main strategy to decrease the prevalence of this virus is vector control (5,12). One potential control strategy is the use of ISVs to target pathogenic viruses through SIE (16–22). The WNV vector Cx. tarsalis is a competent host for the ISV, EILV (13). Therefore, the primary goal of the present study was to determine whether EILV causes SIE against WNV— an unrelated flavivirus virus—in C6/36 cells and Cx. tarsalis mosquitoes. In the present study, we report that EILV suppresses the titers of both WNV strains WN02-1956 and NY99 in C6/36 cells. In contrast, EILV enhanced the viral infection titers of NY99 in Cx. tarsalis mosquitoes at 3 days post superinfection but suppressed WN02-1956 infection titers at 7 days post superinfection. Overall, EILV was found to cause SIE against WNV in both C6/36 cells and Cx. tarsalis mosquitoes but in a strain-specific manner

Previous studies on the interactions between ISVs and other viruses under superinfection conditions have reported variable results, ranging from interference (17,18,22,56,40–42)and enhancement (39,43) to no effect on the secondary infection (38,44). We found that the alphavirus EILV caused SIE against the flavivirus WNV in C6/36 cells, irrespective of the MOI of WNV tested in our study. Superinfection of EILV-infected C6/36 cells with either the WNV strain WN02-1956 or NY99 at an MOI of 0.1 or 0.01 suppressed WNV viral titers as early as 48–72 h post superinfection. Although NY99 titers at an MOI of 0.01 showed signs of recovery towards the final timepoint in our study, these findings are similar to previous findings indicating that ISVs cause heterologous interference in vitro against pathogenic viruses belonging to other families (30,35,45–47)The ability of EILV to cause SIE is also not limited to WNV; it causes SIE against itself and other alphaviruses, including Sindbis virus, western equine encephalitis virus, and CHIKV (22). This indicates that EILV provides a broad range of protection against secondary infection in cells.

The exact mechanism of SIE remains elusive, but interference at different stages of the life cycle of the secondary virus—including attachment (31,32), penetration (24), and replication (33,34)—by the primary virus can contribute to SIE. We showed that EILV significantly reduced the titer of attached NY99 under superinfection conditions in C6/36 cells. The ability of EILV to interfere with NY99 attachment can be explained by downregulation of WNV receptors or co-receptors after EILV infection in these cells, as previously observed regarding other superinfecting viruses (32,48,49). In contrast, the attachment of WN02-1956 was not altered by the presence of EILV in our study. The differences observed between the WNV strains are potentially owing to the mutations in the E and NS5 protein nucleotide sequences of the WN02 genotype compared the nucleotide sequences of these proteins in NY99 genotype (7,8). These mutations may help superinfecting WN02-1956 to escape or compensate for the interference caused by EILV at the attachment stage in C6/36 cells (9); however, further studies are needed to confirm the mechanism of strain-specific interference caused by EILV at the attachment stage. Although the interference at the attachment stage by EILV likely contributes to the mechanism of SIE, it is not the main cause because at a later stage in the viral life cycle—internalization—the titers of both superinfecting strains NY99 and WN02-1956 were similar in the presence or absence of EILV.

The RNAi defense pathway is another potential driver of SIE in mosquitoes and mosquito cell lines (50,51). However, our use of C6/36 cells excluded siRNA, the main antiviral defense pathway, as the root cause of the ability of EILV to suppress WNV titers in these cells because C6/36 cells lack a functional dicer-2 vital for siRNA activation (52). However, the PIWI-interacting RNA (piRNA) pathway remains active in these cells (52,53). A local pairwise comparison of the EILV genome with that of WN02-1956 or NY99 confirmed that these viruses do not share sequence similarities of 21–30 nucleotides in length (not shown), necessary for siRNA and piRNA pathways of cross-protection (52,54). The ability of CHIKV—another alphavirus—to cause heterologous interference was similarly independent of the RNAi pathway (30). Small RNA analysis of EILV and WNV superinfection in C6/36 cells may clarify the role of the RNAi pathway in the ability of EILV to suppress WNV infection in these cells.

Many superinfection studies have been performed in cell culture but not in mosquito vectors. In the present study, we found that EILV did not alter the vector competence of either WNV strain in Cx. tarsalis under superinfection conditions at any timepoint in our study. Although EILV did enhance the NY99 whole-body titers in Cx. tarsalis, the WN02-1956 body titers remained unaffected at 3 days post superinfection. Similarly, a previous study has reported that the ISV Culex flavivirus Izabal enhanced WNV transmission in Cx. quinquefasciatus (Honduras colony) mosquitoes when co-infected (38). A likely explanation for the enhanced NY99 titers in our study is that NY99 “piggybacks” and infects more cells or cell types with the help of EILV. This strategy is used by the human immunodeficiency virus during co-infection with viruses such as Epstein-Barr virus to expand its tissue tropism in its host (55). In the present study, at the later timepoint, 7 days post superinfection—NY99 infection titers were no longer enhanced, whereas WN02-1956 infection titers were suppressed. Previous studies have reported that other ISVs also suppress WNV, but these studies used only one WNV strain to determine SIE (19,38,40). The leg and saliva titers of both NY99 and WN02-1956 were not affected by EILV at this timepoint. NY99 and WN02-1956 are genetically distinct, and this genetic variation may explain the different results obtained in the present study (7–9). The mutations in the WN02 genotype enhance its dissemination in Cx. tarsalis compared with that of NY99 (9). The decrease in WN02-1956 whole-body titers may be attributed to a need for more shared resources with EILV at a faster rate than NY99 (9). Further investigation into the effect of individual mutations between NY99 and WN02-1956 or transcriptional differences between NY99 and WN02-1956 under superinfection conditions with EILV may elucidate the strain-specific effect of EILV on WNV in Cx. tarsalis.

A caveat of our study on EILV–WNV superinfection is the use of an acute EILV infection instead of a persistent infection as observed for most ISVs in nature (16,56,57). ISVs are thought to be vertically transmitted from mother to offspring in nature (40,56,57); however, although EILV infects the ovaries of Cx. tarsalis following an oral challenge, EILV does not exhibit vertical transmission in laboratory conditions (13). A persistent EILV infection may result in different outcomes when followed by superinfection with WNV. Alternatively, the intrathoracic route of infection mimics an ISV infection better than the oral route in mosquitoes (13). However, because EILV infection via the intrathoracic route did not result in the infection of Cx. tarsalis ovaries (13), we used the oral route of infection in the present study to maximize the likelihood of EILV–WNV interaction.

Overall, our study adds to the growing literature on ISVs and their ability to cause SIE against human pathogenic viruses (17,19,37,38,40). ISVs have been reported to be a potentially safe tool to target pathogenic viruses via SIE in nature (54), but our study demonstrates that although EILV caused heterologous interference against both WNV strains—NY99 and WN02-1956—in C6/36 cells, SIE in Cx. tarsalis was strain-specific. The WNV strain NY99 no longer circulates in the US population, but its enhancement in Cx. tarsalis by EILV is a dangerous outcome for a potential tool. Additionally, our results demonstrate the importance of testing all circulating virus strains to determine the safety and ability of an ISV to cause SIE. The suppression of WN02-1956 in Cx. tarsalis was modest, with no effect on the dissemination or transmission of the virus, indicating that SIE alone is inadequate as a control strategy. Paratransgenesis (47,58,59), a strategy by which genetically modified EILV expressing antiviral peptides such as vago (60) is used to target WNV in Cx. tarsalis, may potentially improve the ability of EILV to suppress WNV and increase its specificity.

## ACKNOWLEDGEMENTS

The authors acknowledge Mrs. Melissa James, Mrs. Irene Miller and Mrs. Lindsey LaBella for their assistance in coordinating and managing experiments and facilities at Eva J. Pell lab (University Park, PA, USA). We also acknowledge Mrs. Amelia Romo for her assistance with mosquito rearing, and (University Park, PA, USA) and Dr. Sultan Asad for assistance with experiments. This study was supported by NIH/NIAID grants R01A1128201 and R01AI150251, NSF/BIO grant 1645331, USDA Hatch Project 4769, a grant with the Pennsylvania Department of Health using Tobacco Settlement Funds, and funds from the Dorothy Foehr Huck and J. Lloyd Huck endowment to JLR.

## REFERENCES

1. Knipe DM, Howley PM, Cohen JI, Griffin DE, Lamb RA, Martin MA, et al. Flaviviruses. In: Fields Virology. 6th ed. Lippincott Williams and Wilkins, Baltimore; 2013. p. 747–94.

2. Enis D, Ash N, Arzad F, Ostashari M, Nnie A, Ine F, et al. THE OUTBREAK OF WEST NILE VIRUS INFECTION IN THE NEW YORK CITY AREA IN 1999 A BSTRACT. N Engl J Med [Internet]. 2001 [cited 2023 Feb 21];344(24). Available from: www.nejm.org

3. Sejvar JJ. West Nile Virus: An Historical Overview. Ochsner J [Internet]. 2003 Jun [cited 2022 Oct 12];5(3):6. Available from: /pmc/articles/PMC3111838/

4. Gubler DJ. The continuing spread of West Nile virus in the Western Hemisphere. Vol. 45, Clinical Infectious Diseases. 2007. p. 1039–46.

5. Center of Disease Control and Prevention (CDC). West Nile virus | West Nile Virus | CDC [Internet]. Center of Disease Control and Prevention (CDC). 2019 [cited 2020 Nov 24]. Available from: https://www.cdc.gov/westnile/index.html

6. CDC. Final Cumulative Maps and Data | West Nile Virus | CDC [Internet]. Centers for Disease Control and Prevention. 2020 [cited 2022 Aug 18]. Available from: https://www.cdc.gov/westnile/statsmaps/cumMapsData.html#one

7. Davis CT, Ebel GD, Lanciotti RS, Brault AC, Guzman H, Siirin M, et al. Phylogenetic analysis of North American West Nile virus isolates, 2001-2004: Evidence for the emergence of a dominant genotype. Virology. 2005 Nov 25;342(2):252–65.

8. Snapinn KW, Holmes EC, Young DS, Bernard KA, Kramer LD, Ebel GD. Declining Growth Rate of West Nile Virus in North America. J Virol [Internet]. 2007 Mar [cited 2023 Apr 24];81(5):2531–4. Available from: https://journals.asm.org/doi/10.1128/JVI.02169-06

9. Moudy RM, Meola MA, Morin LLL, Ebel GD, Kramer LD. A newly emergent genotype of West Nile virus is transmitted earlier and more efficiently by Culex mosquitoes. Am J Trop Med Hyg [Internet]. 2007 [cited 2023 Feb 21];77(2):365–70. Available from: https://www.researchgate.net/publication/6150051_A_Newly_Emergent_Genotype_of_West_Nile_Virus_Is_Transmitted_Earlier_and_More_Efficiently_by_Culex_Mosquitoes

10. Dunphy BM, Kovach KB, Gehrke EJ, Field EN, Rowley WA, Bartholomay LC, et al. Long-term surveillance defines spatial and temporal patterns implicating Culex tarsalis as the primary vector of West Nile virus. Sci Reports 2019 91 [Internet]. 2019 Apr 29 [cited 2022 Oct 12];9(1):1–10. Available from: https://www.nature.com/articles/s41598-019-43246-y

11. Deichmeister JM, Telang A. Abundance of West Nile virus mosquito vectors in relation to climate and landscape variables. J Vector Ecol [Internet]. 2011 Jun 1 [cited 2023 Feb 21];36(1):75–85. Available from: https://onlinelibrary.wiley.com/doi/full/10.1111/j.1948-7134.2011.00143.x

12. Gubler DJ, Campbell GL, Nasci R, Komar N, Petersen L, Roehrig JT. West Nile virus in the United States: Guidelines for detection, prevention, and control. Vol. 13, Viral Immunology. Fort Collins; 2000. p. 469–75.

13. Joseph RE, Urakova N, Werling KL, Metz HC, Montanari K, Rasgon JL. Culex tarsalis Is a Competent Host of the Insect-Specific Alphavirus Eilat Virus (EILV). Heise MT, editor. J Virol [Internet]. 2023 Apr 26 [cited 2023 May 10]; Available from: https://journals.asm.org/doi/10.1128/jvi.01960-22

14. Nasar F, Palacios G, Gorchakov R V., Guzman H, Travassos Da Rosa AP, Savji N, et al. Eilat virus, a unique alphavirus with host range restricted to insects by RNA replication. Proc Natl Acad Sci U S A [Internet]. 2012 Sep 4 [cited 2020 Oct 25];109(36):14622–7. Available from: /pmc/articles/PMC3437828/?report=abstract

15. Nasar F, Gorchakov R V., Tesh RB, Weaver SC. Eilat Virus Host Range Restriction Is Present at Multiple Levels of the Virus Life Cycle. J Virol [Internet]. 2015 Jan 15 [cited 2020 Oct 26];89(2):1404–18. Available from: http://jvi.asm.org/

16. Bolling BG, Olea-Popelka FJ, Eisen L, Moore CG, Blair CD. Transmission dynamics of an insect-specific flavivirus in a naturally infected Culex pipiens laboratory colony and effects of co-infection on vector competence for West Nile virus. Virology. 2012 Jun 5;427(2):90–7.

17. Hall-Mendelin S, McLean BJ, Bielefeldt-Ohmann H, Hobson-Peters J, Hall RA, Van Den Hurk AF. The insect-specific Palm Creek virus modulates West Nile virus infection in and transmission by Australian mosquitoes. Parasites and Vectors [Internet]. 2016 Jul 25 [cited 2022 Jun 15];9(1):1–10. Available from: https://link.springer.com/articles/10.1186/s13071-016-1683-2

18. Pauvolid-Corrêa A, Solberg O, Couto-Lima D, Kenney J, Serra-Freire N, Brault A, et al. Nhumirim virus, a novel flavivirus isolated from mosquitoes from the Pantanal, Brazil. Arch Virol [Internet]. 2015 Jan 1 [cited 2023 Apr 15];160(1):21. Available from: /pmc/articles/PMC4785999/

19. Goenaga S, Kenney JL, Duggal NK, Delorey M, Ebel GD, Zhang B, et al. Potential for Co-Infection of a Mosquito-Specific Flavivirus, Nhumirim Virus, to Block West Nile Virus Transmission in Mosquitoes. Viruses [Internet]. 2015 Nov 11 [cited 2023 Apr 15];7(11):5801. Available from: /pmc/articles/PMC4664984/

20. Patterson EI, Kautz TF, Contreras-Gutierrez MA, Guzman H, Tesh RB, Hughes GL, et al. Negeviruses Reduce Replication of Alphaviruses during Coinfection. J Virol [Internet]. 2021 Jun 24 [cited 2023 Mar 13];95(14):433–54. Available from: https://journals.asm.org/doi/10.1128/JVI.00433-21

21. Urakova N, Brustolin M, Joseph RE, Johnson RM, Pujhari S, Rasgon JL. Anopheles gambiae densovirus (AgDNV) negatively affects Mayaro virus infection in Anopheles gambiae cells and mosquitoes. Parasites and Vectors [Internet]. 2020 Apr 22 [cited 2020 Oct 25];13(1):210. Available from: https://parasitesandvectors.biomedcentral.com/articles/10.1186/s13071-020-04072-8

22. Nasar F, Erasmus JH, Haddow AD, Tesh RB, Weaver SC. Eilat virus induces both homologous and heterologous interference. Virology. 2015 Oct 1;484:51–8.

23. Johnston RE, Wan K, Bose HR. Homologous Interference Induced by Sindbis Virus. J Virol [Internet]. 1974 Nov [cited 2022 Oct 13];14(5):1076. Available from: /pmc/articles/PMC355622/?report=abstract

24. Singh IR, Suomalainen M, Varadarajan S, Garoff H, Helenius A. Multiple mechanisms for the inhibition of entry and uncoating of superinfecting Semliki Forest virus. Virology. 1997 Apr 28;231(1):59–71.

25. Zou G, Zhang B, Lim P-Y, Yuan Z, Bernard KA, Shi P-Y. Exclusion of West Nile Virus Superinfection through RNA Replication. J Virol [Internet]. 2009 Nov 15 [cited 2022 Oct 13];83(22):11765. Available from: /pmc/articles/PMC2772679/

26. Eaton BT. Heterologous interference in Aedes albopictus cells infected with alphaviruses. J Virol [Internet]. 1979 Apr [cited 2023 Feb 21];30(1):45–55. Available from: https://journals.asm.org/journal/jvi

27. Karpf AR, Lenches E, Strauss EG, Strauss JH, Brown DT. Superinfection exclusion of alphaviruses in three mosquito cell lines persistently infected with Sindbis virus. J Virol [Internet]. 1997 Sep [cited 2023 Apr 4];71(9):7119–23. Available from: https://journals.asm.org/doi/10.1128/jvi.71.9.7119-7123.1997

28. Abrao EP, Da Fonseca BAL. Infection of Mosquito Cells (C6/36) by Dengue-2 Virus Interferes with Subsequent Infection by Yellow Fever Virus. https://home.liebertpub.com/vbz [Internet]. 2016 Feb 5 [cited 2023 Apr 4];16(2):124–30. Available from: https://www.liebertpub.com/doi/10.1089/vbz.2015.1804

29. Fujita R, Kato F, Kobayashi D, Murota K, Takasaki T, Tajima S, et al. Persistent viruses in mosquito cultured cell line suppress multiplication of flaviviruses. Heliyon. 2018 Aug 1;4(8).

30. Boussier J, Levi L, Weger-Lucarelli J, Poirier EZ, Vignuzzi M, Albert ML. Chikungunya virus superinfection exclusion is mediated by a block in viral replication and does not rely on non-structural protein 2. PLoS One [Internet]. 2020 Nov 1 [cited 2023 Mar 13];15(11). Available from: /pmc/articles/PMC7660575/

31. Wildum S, Schindler M, Münch J, Kirchhoff F. Contribution of Vpu, Env, and Nef to CD4 Down-Modulation and Resistance of Human Immunodeficiency Virus Type 1-Infected T Cells to Superinfection. J Virol [Internet]. 2006 Aug 15 [cited 2023 Apr 4];80(16):8047–59. Available from: https://journals.asm.org/doi/10.1128/JVI.00252-06

32. Walters K-A, Joyce MA, Addison WR, Fischer KP, Tyrrell DLJ. Superinfection Exclusion in Duck Hepatitis B Virus Infection Is Mediated by the Large Surface Antigen. J Virol [Internet]. 2004 Aug [cited 2023 Apr 4];78(15):7925–37. Available from: https://journals.asm.org/doi/10.1128/JVI.78.15.7925-7937.2004

33. Lee Y-M, Tscherne DM, Yun S-I, Frolov I, Rice CM. Dual Mechanisms of Pestiviral Superinfection Exclusion at Entry and RNA Replication. J Virol [Internet]. 2005 Mar 15 [cited 2023 Apr 4];79(6):3231–42. Available from: https://journals.asm.org/doi/10.1128/JVI.79.6.3231-3242.2005

34. Adams RH, Brown DT. BHK cells expressing Sindbis virus-induced homologous interference allow the translation of nonstructural genes of superinfecting virus. J Virol [Internet]. 1985 May [cited 2022 Oct 10];54(2):351–7. Available from: https://journals.asm.org/doi/10.1128/jvi.54.2.351-357.1985

35. Schultz MJ, Frydman HM, Connor JH. Dual Insect specific virus infection limits Arbovirus replication in Aedes mosquito cells. Virology. 2018 May 1;518:406–13.

36. Contreras-Gutierrez MA, Guzman H, Thangamani S, Vasilakis N, Tesh RB. Experimental Infection with and Maintenance of Cell Fusing Agent Virus (Flavivirus) in Aedes aegypti. Am J Trop Med Hyg [Internet]. 2017 Jul 7 [cited 2022 Jun 15];97(1):299. Available from: /pmc/articles/PMC5508911/

37. Hobson-Peters J, Yam AWY, Lu JWF, Setoh YX, May FJ, Kurucz N, et al. A New Insect-Specific Flavivirus from Northern Australia Suppresses Replication of West Nile Virus and Murray Valley Encephalitis Virus in Co-infected Mosquito Cells. PLoS One [Internet]. 2013 Feb 27 [cited 2023 Apr 15];8(2). Available from: /pmc/articles/PMC3584062/

38. Kent RJ, Crabtree MB, Miller BR. Transmission of West Nile Virus by Culex quinquefasciatus Say Infected with Culex Flavivirus Izabal. PLoS Negl Trop Dis [Internet]. 2010 May [cited 2022 Oct 13];4(5):e671. Available from: https://journals.plos.org/plosntds/article?id=10.1371/journal.pntd.0000671

39. Chomczynski P. A reagent for the single-step simultaneous isolation of RNA, DNA and proteins from cell and tissue samples. Biotechniques [Internet]. 1993 [cited 2022 Oct 13];15(3):532–7. Available from: https://pubmed.ncbi.nlm.nih.gov/7692896/

40. Bolling BG, Eisen L, Moore CG, Blair CD. Insect-Specific Flaviviruses from Culex Mosquitoes in Colorado, with Evidence of Vertical Transmission. Am J Trop Med Hyg [Internet]. 2011 Jul 7 [cited 2022 Jun 5];85(1):169. Available from: /pmc/articles/PMC3122363/

41. Colmant AMG, Hobson-Peters J, Bielefeldt-Ohmann H, van den Hurk AF, Hall-Mendelin S, Chow WK, et al. A New Clade of Insect-Specific Flaviviruses from Australian Anopheles Mosquitoes Displays Species-Specific Host Restriction . mSphere [Internet]. 2017 Aug 30 [cited 2022 May 15];2(4). Available from: https://journals.asm.org/doi/full/10.1128/mSphere.00262-17

42. Goenaga S, Goenaga J, Boaglio ER, Alcira Enria D, Del S, Levis C. Superinfection exclusion studies using West Nile virus and Culex flavivirus strains from Argentina.

43. Kuwata R, Isawa H, Hoshino K, Sasaki T, Kobayashi M, Maeda K, et al. Analysis of Mosquito-Borne Flavivirus Superinfection in Culex tritaeniorhynchus (Diptera: Culicidae)

44. Cells Persistently Infected with Culex Flavivirus (Flaviviridae). J Med Entomol [Internet]. 2015 Mar 1 [cited 2023 Apr 24];52(2):222–9. Available from: https://academic.oup.com/jme/article/52/2/222/887669

44. Talavera S, Birnberg L, Nuñez AI, Muñoz-Muñoz F, Vázquez A, Busquets N. Culex flavivirus infection in a Culex pipiens mosquito colony and its effects on vector competence for Rift Valley fever phlebovirus. Parasit Vectors [Internet]. 2018 May 23 [cited 2023 Apr 24];11(1). Available from: /pmc/articles/PMC5966921/

45. Fujita R, Kato F, Kobayashi D, Murota K, Takasaki T, Tajima S, et al. Persistent viruses in mosquito cultured cell line suppress multiplication of flaviviruses. Heliyon. 2018 Aug 1;4(8):e00736.

46. Urakova N, Brustolin M, Joseph RE, Johnson RM, Pujhari S, Rasgon JL. Anopheles gambiae densovirus (AgDNV) negatively affects Mayaro virus infection in Anopheles gambiae cells and mosquitoes. Parasites and Vectors [Internet]. 2020 Apr 22 [cited 2020 Oct 29];13(1):210. Available from: https://parasitesandvectors.biomedcentral.com/articles/10.1186/s13071-020-04072-8

47. Patterson EI, Kautz TF, Contreras-Gutierrez MA, Guzman H, Tesh RB, Hughes GL, et al. Negeviruses Reduce Replication of Alphaviruses during Coinfection. J Virol [Internet]. 2021 Jun 24 [cited 2022 Oct 20];95(14):433–54. Available from: /pmc/articles/PMC8223947/

48. Huang I-C, Li W, Sui J, Marasco W, Choe H, Farzan M. Influenza A Virus Neuraminidase Limits Viral Superinfection. J Virol [Internet]. 2008 May 15 [cited 2023 Apr 25];82(10):4834–43. Available from: http://www.fda.gov/cder

49. Michel N, Allespach I, Venzke S, Fackler OT, Keppler OT. The nef protein of human immunodeficiency virus establishes superinfection immunity by a dual strategy to downregulate cell-surface CCR5 and CD4. Curr Biol. 2005 Apr 26;15(8):714–23.

50. Gusachenko ON, Woodford L, Balbirnie-Cumming K, Evans DJ. First come, first served: superinfection exclusion in Deformed wing virus is dependent upon sequence identity and not the order of virus acquisition. ISME J 2021 1512 [Internet]. 2021 Jun 30 [cited 2023 Apr 25];15(12):3704–13. Available from: https://www.nature.com/articles/s41396-021-01043-4

51. Ratcliff FG, MacFarlane SA, Baulcombe DC. Gene Silencing without DNA: RNA-Mediated Cross-Protection between Viruses. Plant Cell [Internet]. 1999 Jul 1 [cited 2023 Apr 25];11(7):1207–15. Available from: https://academic.oup.com/plcell/article/11/7/1207/6008433

52. Brackney DE, Scott JC, Sagawa F, Woodward JE, Miller NA, Schilkey FD, et al. C6/36 Aedes albopictus Cells Have a Dysfunctional Antiviral RNA Interference Response. PLoS Negl Trop Dis [Internet]. 2010 Oct [cited 2023 Apr 25];4(10):856. Available from: /pmc/articles/PMC2964293/

53. Varjak M, Leggewie M, Schnettler E. The antiviral piRNA response in mosquitoes? J Gen Virol [Internet]. 2018 Dec 1 [cited 2023 Apr 25];99(12):1551–62. Available from: https://www.microbiologyresearch.org/content/journal/jgv/10.1099/jgv.0.001157/sidebyside

54. Laureti M, Paradkar PN, Fazakerley JK, Rodriguez-Andres J. Superinfection exclusion in mosquitoes and its potential as an arbovirus control strategy [Internet]. Vol. 12, Viruses. 2020. Available from: www.mdpi.com/journal/viruses

55. Root-Bernstein RS, Hobbs SH. Does HIV “Piggyback” on CD4-like Surface Proteins of Sperm, Viruses, and Bacteria? Implications for Co-transmission, Cellular Tropism and the Induction of Autoimmunity in AIDS. J Theor Biol. 1993 Jan 21;160(2):249–64.

56. Saiyasombat R, Bolling BG, Brault AC, Bartholomay LC, Blitvich BJ. Evidence of Efficient Transovarial Transmission of Culex Flavivirus by Culex pipiens (Diptera: Culicidae). J Med Entomol [Internet]. 2011 Sep 1 [cited 2022 Jun 5];48(5):1031–8. Available from: https://academic.oup.com/jme/article/48/5/1031/910470

57. Duggal NK, Langwig KE, Ebel GD, Brault AC. On the Fly: Interactions Between Birds, Mosquitoes, and Environment That Have Molded West Nile Virus Genomic Structure Over Two Decades. J Med Entomol [Internet]. 2019 Oct 28 [cited 2022 Jul 1];56(6):1467– 74. Available from: https://academic.oup.com/jme/article/56/6/1467/5572129

58. Ren X, Hoiczyk E, Rasgon JL. Viral Paratransgenesis in the Malaria Vector Anopheles gambiae. Schneider DS, editor. PLoS Pathog [Internet]. 2008 Aug 22 [cited 2020 Nov 25];4(8):e1000135. Available from: https://dx.plos.org/10.1371/journal.ppat.1000135

59. Adelman ZN, Blair CD, Carlson JO, Beaty BJ, Olson KE. Sindbis virus-induced silencing of dengue viruses in mosquitoes. Insect Mol Biol [Internet]. 2001 Jun 1 [cited 2020 Oct 26];10(3):265–73. Available from: https://www.onlinelibrary.wiley.com/doi/full/10.1046/j.1365-2583.2001.00267.x

60. Paradkar PN, Trinidad L, Voysey R, Duchemin JB, Walker PJ. Secreted Vago restricts West Nile virus infection in Culex mosquito cells by activating the Jak-STAT pathway. Proc Natl Acad Sci U S A [Internet]. 2012 Nov 13 [cited 2020 Nov 24];109(46):18915–20. Available from: www.pnas.org/cgi/doi/10.1073/pnas.1205231109

